# High-throughput UHPLC-MS to screen metabolites in feces for gut metabolic health

**DOI:** 10.1101/2021.12.22.473790

**Authors:** Andressa de Zawadzki, Maja Thiele, Tommi Suvitaival, Asger Wretlind, Min Kim, Mina Ali, Annette F. Bjerre, Karin Stahr, Ismo Matilla, Torben Hansen, Aleksander Krag, Cristina Legido-Quigley

**Affiliations:** Steno Diabetes Center Copenhagen, Herlev, Denmark; Department of Gastroenterology and Hepatology, Odense University Hospital, Odense, Denmark; Department of Clinical Medicine, University of Southern Denmark, Odense, Denmark; Copenhagen Prospective Studies on Asthma in Childhood, Herlev and Gentofte Hospital, Denmark; Novo Nordisk Foundation Center for Basic Metabolic Research, Faculty of Health and Medical Sciences, University of Copenhagen, Copenhagen, Denmark; Institute of Pharmaceutical Science, King’s College London, London, UK

**Author notes:** **Corresponding Author**, Cristina Legido-Quigley -.

**Keywords:** bile acids, fecal metabolomics, gut-liver axis, targeted metabolomics, sample preparation

## Abstract

**Background:** Feces are the product of our diets and have been linked to diseases of the gut, including Chron’s disease and metabolic diseases such as diabetes. For screening metabolites in heterogeneous samples such as feces, it is necessary to use fast and reproducible analytical methods that maximize metabolite detection.

**Methods:** As sample preparation is crucial to obtain high quality data in MS-based clinical metabolomics, we developed a novel, efficient and robust method for preparing fecal samples for analysis with a focus in reducing aliquoting and detecting both polar and non-polar metabolites. Fecal samples (n= 475) from patients with alcohol-related liver disease and healthy controls were prepared according to the proposed method and analyzed in an UHPLC-QQQ targeted platform in order to obtain a quantitative profile of compounds that impact liver-gut axis metabolism.

**Results:** MS analyses of the prepared fecal samples have shown reproducibility and coverage of n=28 metabolites, mostly comprising bile acids and amino acids. We report metabolite-wise relative standard deviation (RSD) in quality control samples, inter-day repeatability, LOD, LOQ and range of linearity. The average concen-trations for 135 healthy participants are reported here for clinical applications.

**Conclusions:** our high-throughput method provides an efficient tool for investigating gut-liver axis metabolism in liver-related diseases using a noninvasive collected sample.

## 1. INTRODUCTION

Sample preparation is a critical step for obtaining high quality data in MS-based methods. An ideal sample preparation method requires a fast and reproducible method that allows to screen a large variety of metabolites and maximizes metabolite detection ^1,2^. High standards of method reproducibility are especially important in clinical studies, as they usually involve a large number of biological samples from various matrices such as plasma, urine, tissue and feces ^1,3,4^. Due to the heterogeneity of fecal samples, there is a need for standardized protocols of sample preparation for analysis that allows direct comparisons between cohorts ^2,5–7^.

The molecular makeup of feces has gained increasing interest from clinical researchers, as collecting samples is non-invasive and the fecal metabolome provides a readout of the host-gut microbiota interactions and of the function of other organs in close contact with gut activity ^2,8,9^like the liver ^10,11^. Among the small molecules detected in feces that regulate host-microbiota interactions and liver activity, bile acids have been widely reported as potential biomarkers for liver diseases, metabolic syndrome and dysbiosis ^8,12–14^. Amino acids from the diet are also redirected to important metabolic pathways such as bile acid synthesis ^15^ by gut microbiota. Several studies, including ours, have demonstrated that metabolic signatures of amino acids can assist the prediction and diagnosis of diseases like type 2 diabetes ^16^, fatty liver diseases ^17,18^, metabolic syndrome and obesity ^19^, depression and neurophysiological diseases ^20^. Finally, other compounds excreted in feces, such as kynurenine ^21^ and azelaic acid ^22^, are involved in the pathogenesis of metabolic syndrome and liver diseases.

In general sample preparation methods for fecal samples focus on one class of compounds at a time due to the complex matrix. The quantification of bile acids in feces has been successfully applied to ^7 14 15 23 26^ clinical studies for gut-liver health^7,14,15,23–26^, with few studies combining other molecular groups with quantification of bile acids ^27,28^.

The vast literature of analytical methods for simultaneous analysis of diverse compound classes mostly consists of untargeted approaches. Targeted analysis with a broad coverage of compounds and absolute concentrations are a prerequisite for mechanistic investigation of diseases and development of clinical tests for diagnostics ^26,29^. Xie et al., 2021 developed a high-throughput metabolite array technology for quantitative determination of up to 300 metabolites for precision medicine. In this study, quantitative profiles of fatty acids, amino acids, carbohydrates and bile acids were successfully obtained and 60 samples were used for feces ^26^.

We previously developed an untargeted method that provides a wide coverage of fecal metabolites including amino acids, lipids (diacylglycerols, triacylglycerols and ceramides), fatty acid derivatives, carboxylic acids and phenolic compounds, and demonstrated that a small proportion of metabolites had acceptable analytical variation^5^. This study prompted the development of a quantitative method for feces screening in our clinical setting.

Herein, we propose a novel robust method for preparing fecal samples with focus on reducing aliquoting and aiming to stratify patients in the clinic. We present the results from a UHPLC-MS-based targeted platform for the simultaneous quantification of bile acids, amino acids and other compounds of interest for the gut-liver axis in a cohort of 475 samples.

## 2. EXPERIMENTAL SECTION

### 2.1. Patients

This study used a total of 475 stool samples from a cross-sectional cohort of persons with a history of harmful drinking and matched healthy controls, as described in a previously published study ^5^. Participants were recruited by Odense University Hospital and informed consent was obtained for all subjects prior to inclusion. The cohort was approved by the ethics committee for Region of Southern Denmark (ethical ID S-20120071, S-20160021, S-20170087 and ID S-20160006G; data protection agency 16/3492). All methods involving participants were performed in agreement to the ethical principles of the Declaration of Helsinki.

### 2.2. Chemicals

Reagent grade potassium carbonate (K_2_CO_3_), potassium bicarbonate (KHCO_3_), sodium hydroxide (NaOH) and hydrochloric acid (HCl) were obtained from Sigma-Aldrich (Steinheim, Germany). Analytical grade formic acid (HCOOH) and LC-MS grade isopropanol (IPA), acetonitrile (ACN) and water (H_2_O) were purchased from Fisher Scientific (Fairlawn, NJ, USA). LC-MS grade methanol (MeOH) was purchased from Honeywell International Inc. (Morristown, NJ, USA). HPLC grade dichloromethane (DCM), anhydrous ACN, and 6-aminoquinoline-N-hydroxy-succinimidyl carbamate (AQC) for derivatization of amino acids were purchased from Santa Cruz Biotechnology, Inc. (Dallas, TX, USA). Amino acids, bile acids and the other analytes were obtained from Sigma-Aldrich (Steinheim, Germany); SCB: Santa Cruz Biotechnology, Inc. (Dallas, TX, USA) or; CIL: Cambridge Isotope Laboratories Inc. (Tewksbury, MA, USA), as listed in the reference ^16^. Bile acids lithocholic acid (LCA), glycolithocholic acid (GLCA) and ursodeoxycholic acid (UDCA) were obtained from Cambridge Isotope Laboratories Inc.

### 2.3. Preparation of standards and calibration standards

All analytes and internal standards for UHPLC analysis are listed in Supporting Information (SI) Table S1 and 2. For each analyte and internal standard, stock solutions at concentration of 5.0 mg mL^−1^ were prepared by dissolving the compound in appropriate solution according to solubility: aqueous solution containing 0.1 M HCl, H_2_O:MeOH (90:10, v/v) or in MeOH ^16^. An internal standard mixture (ISTD MIX) was prepared by diluting each one of the internal standard stock solutions to a final solution of 0.45 M carbonate buffer pH 8.9. A solution of 1 M NaOH (3:1, v/v) was used to adjust the pH to the level required for derivatization of amino acids. The final internal standard solution consisted of a set of 31 stable heavy-labelled compounds ^16^ together with three additional internal standards associated to gut-liver axis (LCA, GLCA and UDCA), see Table S1 and S2. In order to construct calibration curves for quantitative analysis, a stock solution containing the 34 non-labeled analytes at concentration of 50.0μg mL^-1^ was prepared by adding 125.0 μL of each analyte at concentration of 5 mg mL^-1^ into 5500 μL of ACN to produce 10 mL of final solution. A dilution series was prepared by further diluting this stock solution at concentration of 50.0 μg mL^-1^ in ACN in order to construct a calibration curve with the following levels: 1.25; 5.0; 10.0; 25.0; 50.0; 75.0; 100.0; 250; 500; 750; 1000; 2500; 5000; 7500; 10,000; 12,500; 25,000 and 50,000 ng mL^-1^. For derivatization of amino acids and related metabolites, a solution of AQC-reagent at concentration of 5 mg mL^−1^ was prepared by dissolving the compound in anhydrous ACN at 55 °C. Blanks for MS analysis consisted of 40 μL of methanol/water 1:1 (v/v) and 20 μL of AQC dissolved in ACN.

### 2.4. Instrumentation

Samples were analyzed using an ultra-high-performance liquid chromatography system (UHPLC) coupled with a triple quadruple mass spectrometer, both, from Agilent Technologies. The 1290 Infinity UHPLC system (Agilent Technologies, Santa Clara, CA, USA) consisted of a binary pump (model G4220A) with a two-channel solvent degasser, a temperature-controlled column compartment (model G1316C), a multi-sampler equipped with a cooler thermostat (model G7167B), a diode-array detector equipped with an Agilent Max-light cartridge cell and a deuterium lamp. The multi-sampler was maintained at 5 ^°^C and set to use two mixtures for cleaning the needle and the needle seat for 8 seconds after each injection: ACN:MeOH:IPA:H_2_O (1:1:1:1, v/v/v/v) containing 0.1% formic acid and 10% DCM in MeOH. The column used for separation was a Kinetex^®^ F5 100 mm × 2.1 mm, packed with 1.7-μm particles (Phenomenex, Torrance, CA, USA). The column was maintained at 40 °C at a flow rate of 0.4 mL min^-1^ using two mobile phases for gradient elution. The mobile phases used for separation consisted of “A” H_2_O + 0.1 % HCOOH and “B” ACN:IPA (2:1, v/v) + 0.1% HCOOH. The following gradient was used: from 0 to 1 min 1% B, from 1 to 1.8 min 1–18% B, from 1.8 to 3.4 min 18–21% B, from 3.4 to 7 min 21–65% B, from 7 to 7.1 min 65–100% B, from 7.1 to 8.9 min 100% B, from 8.9 to 9.00 min 100-99% B. A period of 2.5 min returning to the initial conditions (1% B) was used for column re-equilibration.

Following UHPLC separation, mass spectrometry analysis of samples was conducted on an Agilent 6460 triple quadrupole mass spectrometer (Agilent Technologies, Santa Clara, CA, USA) equipped with an Agilent Jet Stream electrospray ionization source. A Genius 3010 nitrogen generator from PEAK Scientific Instruments Ltd. (Inchinnan, Scotland, UK) was used to produced nitrogen as nebulizing gas (pressure of 45 psi, 300 °C, 5 L min^-1^) and as sheath gas (250 °C, 11 Lmin-1). All data were recorded with Agilent Mass Hunter LC/MS Data Acquisition Software version B.08.02 (Agilent Technologies). Agilent 6460 triple quadrupole mass spectrometer was calibrated with ESI tuning solution from Agilent Technology prior to the analyses.

For MS quantitative analysis, a method based on dynamic multiple reaction monitoring (MRM) mode was set to include all transitions with optimized parameters for ionizing the analytes of interest. An MRM method previously developed in our group for analysis of plasma ^16^ was adapted in order to include three additional bile acids (GLCA, LCA, UDCA) and to correct for matrix effect. All optimization steps were then combined to a final MRM method that allows to ionize all 34 analytes in positive or negative ion modes according their properties, see supporting information Tables S1 and S2. SRM and MRM optimizations were carried out in MassHunter Optimizer software version B.07.00.

### 2.5. Sample preparation

Sample preparation method was designed by combining two different methods developed by our group and previously described in the literature ^5,16^. Extraction of metabolites was adapted from Trošt et al., 2020 and aimed to obtain a main solution containing a wide range of compounds in appropriate concentration for analysis through various MS platforms. Accordingly, 30 mg of sample were aliquoted in Eppendorf tubes and homogenized in 400 mL of methanol for 5 min at 25 Hz with a sample disruptor Qiagen TissueLyser II Laboratory Mixer (Qiagen, Valencia, CA, USA). The homogenized mixture was centrifuged for 10 minutes at 10000g (4 °C). 250 mL of upper phase were collected and transferred to another clean 1.5 mL Eppendorf tube. The supernatant provided a homogeneous methanolic solution containing the extracted metabolites. The supernatant was then dried using Biotage TurboVap^®^ N_2_ dryer (flow rate of 2 bar) for 3-5 hours in order to increase the concentration of metabolites to a level suitable for MS analysis. After drying, the pellet was resuspended in 150 mL MeOH:H_2_O 1:1(v/v) and the resuspended sample was centrifuged for 3 minutes at 10.000g (4 °C). The following steps were adapted from a protocol used for analysis of bile acids in plasma ^16^. 20 mL of the final fecal supernatant were collected and 20 μL ISTDmix was added. The solution spiked with internal standards was further derivatized with AQC as described by Ahonen et al., 2019 ^16^. After preparing the samples, the remaining supernatant was used for preparing pooled samples for quality control (QC) and the leftover was stored at −80 °C for further analyses. For this study, pooled samples were produced by mixing together all the 475 fecal supernatants produced in the step of derivatization. The resulting pooled supernatants were divided in aliquots of 20 μL which were prepared individually and identically to the original samples by adding 20 μL of ISTDmix following derivatization with AQC.

### 2.6. Study design for evaluation of analytical performance and suitability

The reproducibility and feasibility of our method was investigated in a targeted platform based on UHPLC-MS analysis using clinical samples from a cohort of 350 participants with history of harmful drinking and 125 healthy controls. Experiments were designed to include pooled samples, blanks and calibration curves for quality control and evaluation of analytical performance, see Figure S1.

The limit of detection (LOD) and the limit of quantification (LOQ) were determined for each analyte and internal standard in order to evaluate quantitative performance of compounds in fecal samples. The analytical performance and method reproducibility were evaluated through the variation of individual metabolite concentrations in pooled samples in relation to study samples as estimated by relative standard deviation (RSD) values. In addition to the estimation of inter-variation of pooled samples, the analysis of pools was repeated following 5 and 10 days in order to estimate the intra-sample variation.

### 2.7. Data analysis and statistics

The acquired data was pre-processed using Agilent Mass Hunter Quantitative Analysis software (version B.07.00). Calibration curves were constructed for the concentration ranges between 2.5-50,000 μg mL^−1^ for each metabolite. The peak area of each analyte was normalized by its internal standard and then this ratio, named response, was plotted against concentrations. Calibration curves were obtained by linear regression of the normalized peak areas versus concentrations with three points representing of each concentration. The range of linearity of the calibration curves was determined from lower and upper limits of quantification (LLOQ and ULOQ) by considering only the concentrations that produce a linear curve with R-squared values higher than 0.95. LOD and LOQ were estimated based on the standard deviation (SD) of the intercept (σ) and the slope (s) of the calibration curve ^30^. The concentration of the analytes in the study and pooled samples was determined by inverse-regressing the responses using the calibration curves. The resulting data was then processed using an in-house pipeline created with the software “R”. Calculations were carried out to obtain the relative standard deviation (RSD) values from QC samples before and after batch correction for each metabolite. Principal component analysis (PCA) plots and violin plots were created using the ggplot2 package in R ^31^ and used for data visualization.

## 3. RESULTS AND DISCUSSION

### 3.1. Sample preparation

In this study, we aimed to develop a high-throughput analytical method for fecal samples ^5,16^. Our previous untargeted metabolomics work reported the difficulties when extracting metabolites from feces^5^, particularly for metabolites found in plasma relevant to diabetes, liver and kidney diseases ^16^. Here, we report and discuss the steps for preparing fecal samples for targeted analysis of metabolites related to the gut-liver axis. This proposed sample preparation method is illustrated in detail in Figure 1.

**Figure 1.**
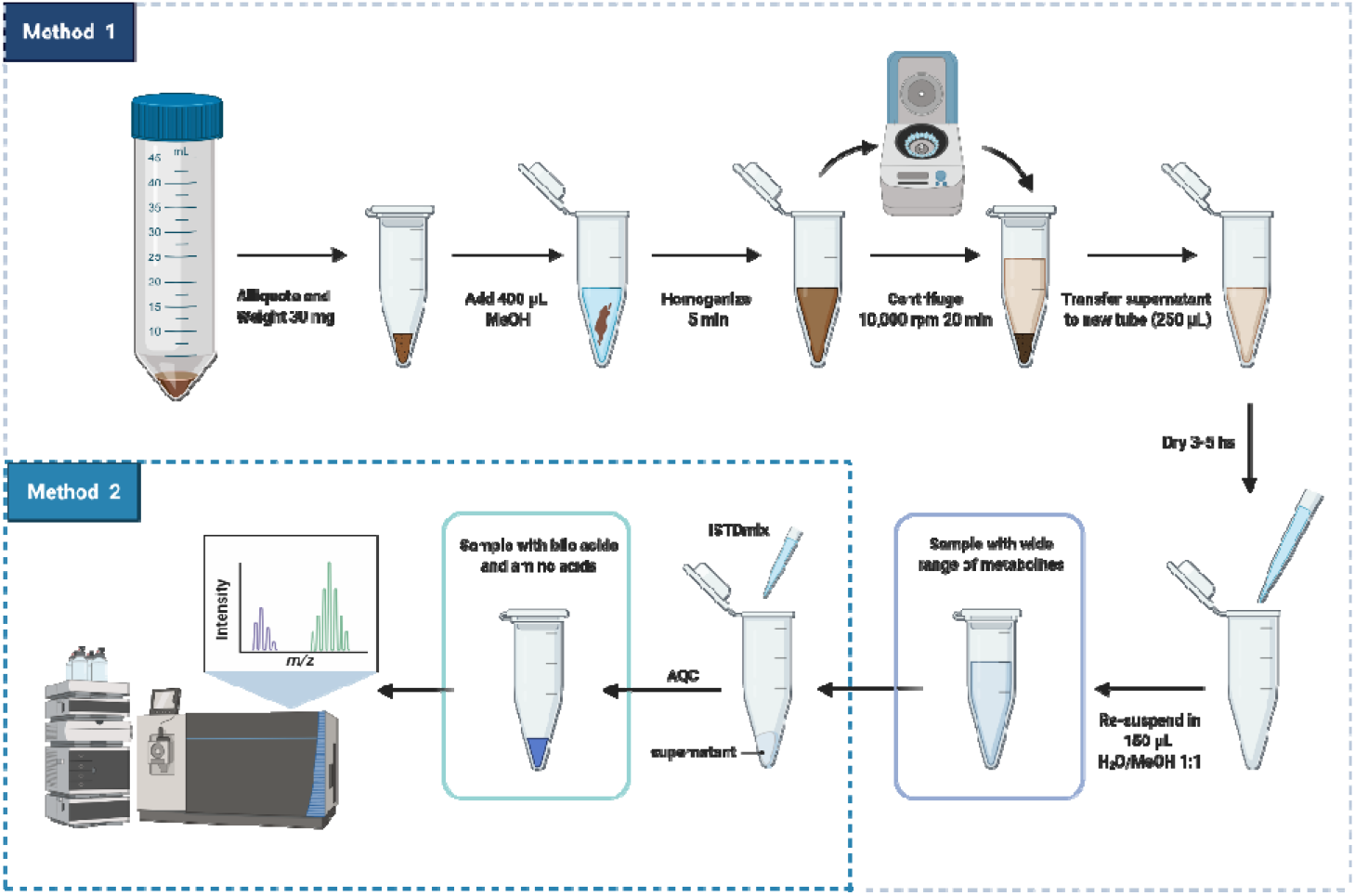
Scheme of sample preparation of fecal samples for targeted analysis of metabolites related to the gut-liver axis. Created with biorender.

As mentioned, in our previous study, Trošt et al., 2020, the metabolite composition of frozen stools from healthy participants presented significant variation between sampling areas due to the heterogeneity of the sample ^5^. For semi-solid complex matrices like feces, it is important that the samples are homogenized prior to metabolite extraction. In the present study, fecal samples undergo three stages of homogenization to correct fluctuations in concentration due to heterogeneity: (i) before being cryogenically drilled and aliquoted to 200 mg, (ii) before being aliquoted to the amount of 30 mg that were used for metabolite extraction and (iii) during metabolite extraction with methanol using bead homogenization.

Methanol was chosen as the solvent of extraction of metabolites from feces due to its versatility for dissolving compounds with different polarities ^1,6^. Methanol has been widely used as a solvent for the extraction of free and conjugated bile acids from human feces for quantitative MS-analysis ^7,23,25^. Methanol also assists the simultaneous extraction of amino acids ^32^, short-chain fatty acids (SCFAs) ^33^ and small organic acids. Additionally, methanol promotes protein denaturation and liberation of protein-bound SCFAs to the organic solvent as well as sample clean-up following a step of centrifugation ^34^.

Our extraction method allowed to obtain several metabolite classes in physiological concentrations to be measured by multiple MS platforms (supporting information, Table S3) ^35^. Besides having bile acids and amino acids, the fecal extract obtained using our method has free fatty acids, carbohydrates and metabolites from tricarboxylic acid cycle (TCA cycle), see Table S3 (supporting information). It is important to highlight that the resultant solution must be subjected to few additional steps of sample preparation before the targeted or untargeted MS analysis of polar metabolites, SCFA, bile acids, amino acids, organic acids and lipids. Herein, we focus the discussion in the reproducibility of the sample preparation method that was developed for targeted analysis of metabolites. For quantification of the compounds, crucial steps, which we will detail next, were added to the core sample preparation method:(i) addition of internal references and (ii) derivatization amino acids into less polar adducts.

#### (i) Addition of internal reference standards

Heavy-labelled isotopes have been widely used as internal references for mass spectroscopy quantitative analysis to ensure system stability and reproducibility. Spiking samples with specific concentration of heavy labelled internal standards identical to the targeted compounds is the most efficient strategy for obtaining absolute concentrations in studies involving large cohorts ^3,36,37^. Moreover, analytical properties of the isotopically labelled reference (m/z, retention time, peak area and peak shape) are efficiently used to correct systematic analytical variations such as drop in MS sensitivity and shifts in retention time ^2,3^. Aiming to apply our method in quantitative studies of large cohorts, we included addition of internal standards to our sample preparation method.

The concentrations of the heavy labelled references are an important parameter to ensure reproducibility. In the present study, concentrations of internal standards were modified from Ahonen et al., 2019 ^16^ to fit the fecal matrix considering the limits of quantification for each metabolite, see supporting information Table S2.

#### (ii) Derivatization of amino acids with AQC reagent

Chromatographic separation of free amino acids is difficult as their similar polarities make them elute through the column at similar retention times. The quantification of free amino acids using liquid chromatographic tandem mass spectrometry usually requires a previous step of derivatization. Derivatization yields adducts with different polarities allowing separation by gradient elution ^38,39^. One advantage of derivatizing amino acids before chromatographic separation is being able to simultaneously analyze other types of compounds. Derivatization of amino acids was included in the sample preparation method in order to allow the simultaneous analysis of bile acids and amino acids in a single chromatographic run ^16^. Using AQC as a derivatization agent has the advantage of fast speed of formation of derivatives ^39,40^. Excess of AQC was used to ensure that the derivatization reaction occurred within few seconds.

### 3.2. Method Optimization

The optimized UHPLC-MS method was validated for analyte separation, limit of detection (LOD), limit of quantification (LOQ), linearity (R_2_) and range of linearity, see Table 1 and supporting information.

**Table 1.**
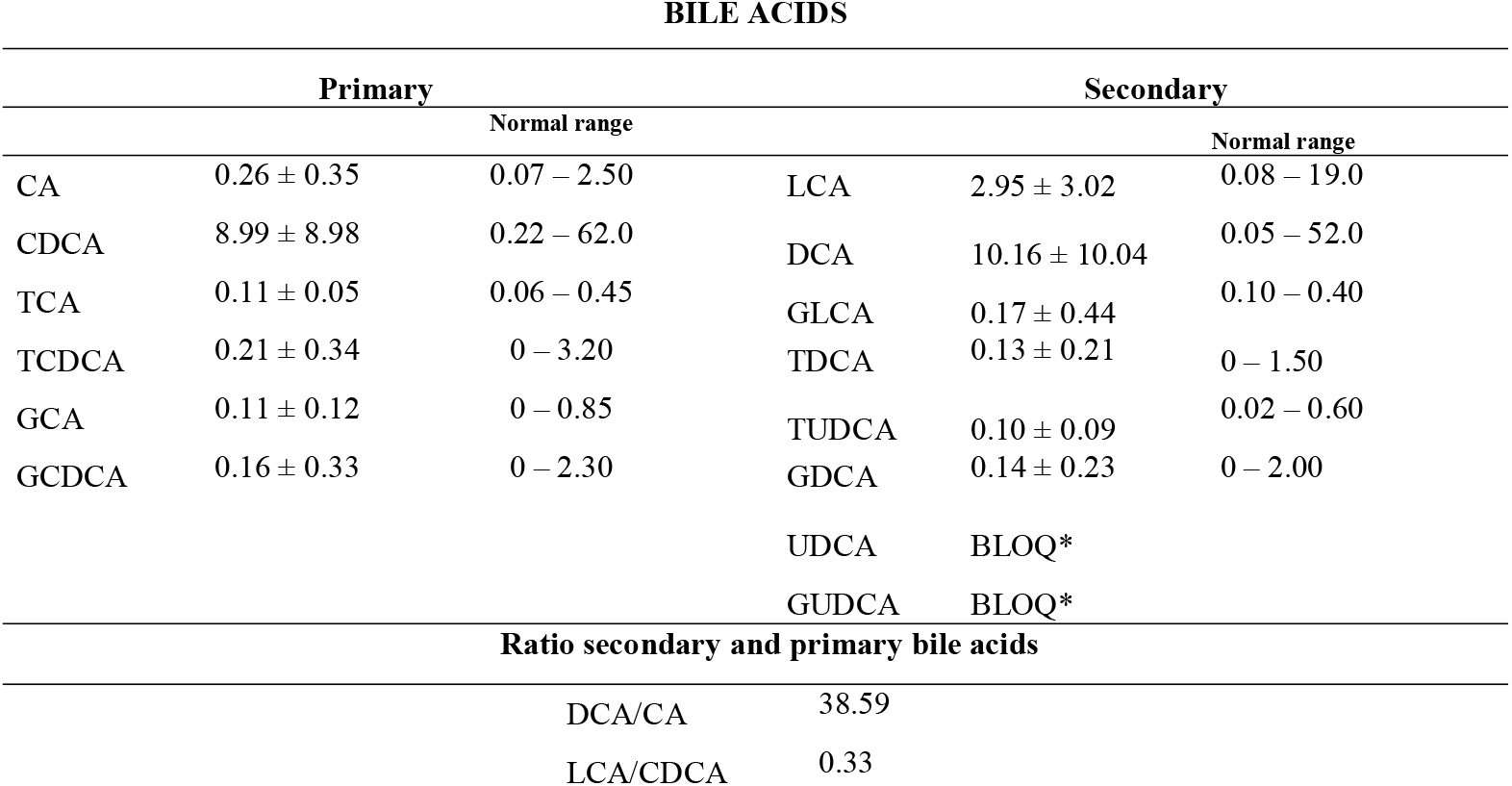
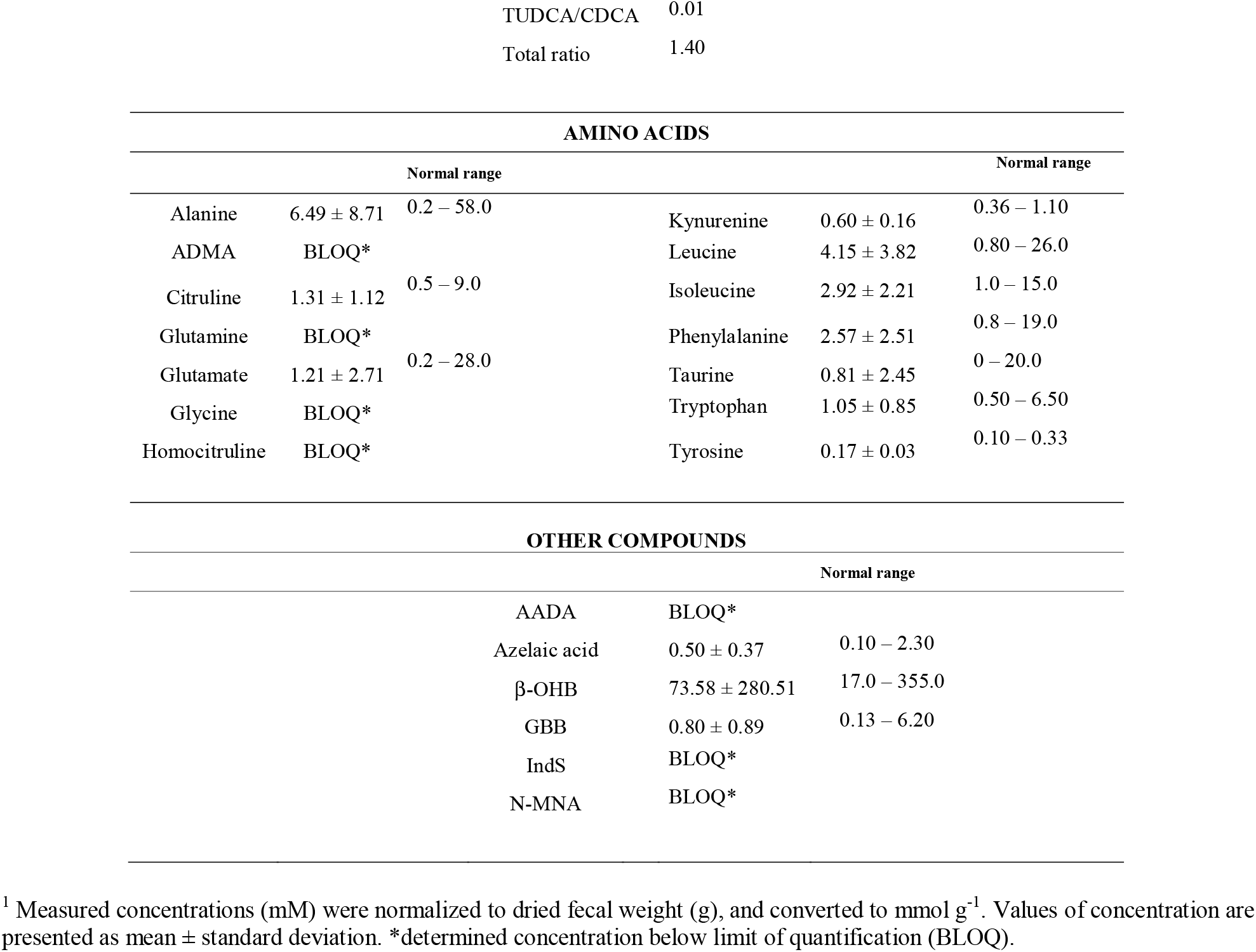
Average concentrations (mmol g^-1^ of dried sample) and range of concentrations of metabolites of relevance for the gut-liver axis in fecal samples from healthy controls.

Fecal extracts spiked with the internal standards presented chromatographic peaks with retention times spanning 0.7-8.0 min (supporting information, Table S1 and S2). The adapted UHPLC-MS method provided good chromatographic separation, sensitivity and selectivity for the determination of 34 analytes investigated. The method developed for the analysis of fecal samples presented retention times very similar to the method that was previously developed for plasma ^16^. However, the new method allowed better separation of the peaks assigned to the pairs Leu/Ile, TDCA/TCDA, GDCA/GCDCA. The improvement was due to the optimization of the MRM method. Additionally, the present method was also optimized to include the quantification of LCA, GLCA and UDCA due to their importance for the gut-liver-axis. LCA, GLCA and UDCA are secondary bile acids produced in the gut by resident microbiota and are associated with protection of gut barrier against inflammation^41,42^. GLCA/GLCA-d4, LCA/LCA-d4 and UDCA/UDCA-d4 exhibited chromatographic peaks at 7.11, 7.57 and 6.9 min, respectively, they were quantified through MRM transitions of 432.3–74.1 / 436.3–74.1, 375.3–375.3 / 379.3–379.3 and 391.3–391.3 / 395.3–395.3, see Tables S1 and S2 in supporting information. Compound identity was validated through retention times and MRM transitions. Then, the limit of detection and the limit of quantification were obtained for each targeted analyte.

The determination of the lowest detectable and quantifiable levels of an analyte, named LOD and LOQ respectively, is essential for method validation as they allow to distinguish analytical properties of analytes from the background ^3^. Analytes showed a wide range of LOD values and the metabolites Ala, CDCA, GCDCA, Leu and TCDCA presented LOD of less than 5 ng mL^−1^. Most of LOD and LOQ for the quantification of bile acids presented in Table S4 (SI) were comparable to the values obtained by Ahonen et al., 2019 ^16^. However, significant differences were found in the quantifiability of amino acids, explained by the change of matrix from plasma (carbonate buffer) to fecal (MeOH:H_2_O 1:1) as well as by the method used to estimating the limits.

For estimation of the quantitative performance, the linear range in which it is possible to quantify the analytes with more than 95% of precision was obtained for each metabolite, see Table S4. LOQ was used for setting the minimum concentration required for construction of the calibration curves (LLOQ). In addition to the determination of LLOQ, the upper limit of quantification was obtained as the highest concentration of analyte that provides a reproducible response with a coefficient of variation (CV) less than 15%. Upper limits of quantification for bile acids were increased in comparison to plasma in order to fit the fecal matrix. The concentration of bile acids in plasma is very low due to an efficient retention of circulating bile acids by the liver. On the other hand, most part of the bile acids produced in the liver and in the gut is excreted in feces where they serve as emulsifiers ^13^. Considering the range of linearity, more than half of analytes exhibited coefficients of determination (R^2^) above 0.99 while most part of the remaining ones showed values above 0.95. Only few analytes presented R_2_ values below 0.95, but they were still within the required accuracy range of 80-120%.

### 3.3. Method feasibility in large/scale cohorts

The reproducibility of the method was investigated considering system stability, analytical performance, precision and feasibility for analysis of a large sample set. Many cohorts used in clinical research comprise large scale studies that require several days for sample MS analysis, which presents unique challenges.

A common strategy for quality assessment in large cohorts is to include QC pooled samples representing replicates throughout the analysis and monitor the behavior of these QC samples over time ^3,43,44^. Pooled quality control samples were added throughout approximately 600 injections to assess the analytical stability of the platform, see Figure 2 and supporting information Figure S2. The measured signal response of each metabolite plotted against the number of injections indicated that the metabolite signal response of the pooled samples presented very little time-related systematic variation. This behavior is demonstrated in Figure 2 through six metabolites: gamma-butyl butyrate (GBB), tryptophan (Trp), leucine (Leu), cholic acid (CA), taurochenoxycholic acid (TCDCA) and deoxycholic acid (DCA). The low variation becomes increasingly evident when QC pooled samples are compared to the samples representing the cohort that presented a wide range of responses, thus indicating method robustness, system stability and good analytical performance over several days for large studies.

**Figure 2.**
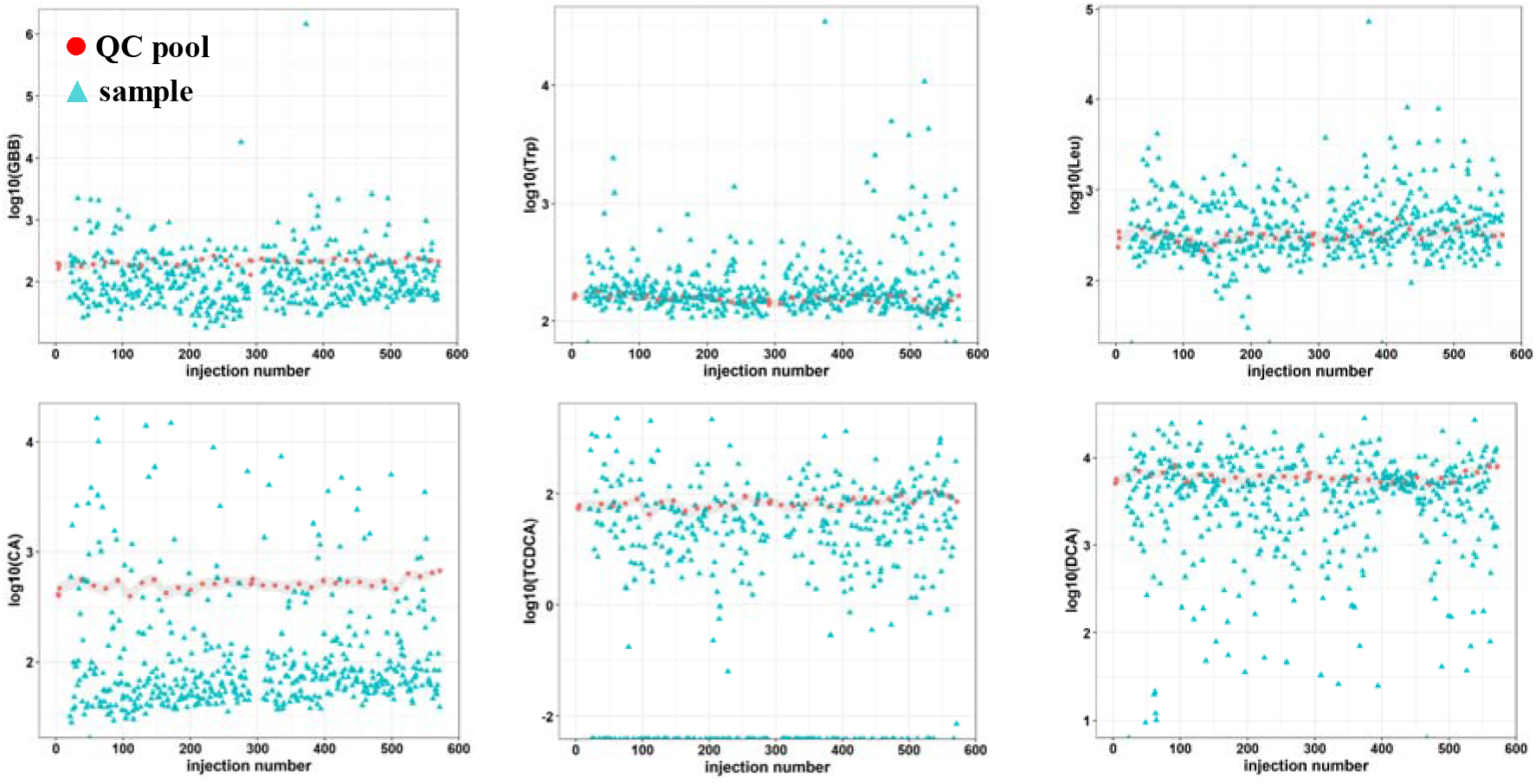
Analytical stability of a targeted UHPLC-MS platform for quantitative analysis of metabolites related to gut-liver axis: gamma-butyl butyrate (GBB), tryptophan (Trp), leucine (Leu), cholic acid (CA), taurochendeoxycholic acid (TCDCA) and deoxycholic acid (DCA). Time related variation of metabolite response measured in quality control pooled samples representing replicates and samples representing a cohort. Interval of variation in QC pooled samples is represented in grey.

Additionally, inter-day repeatability was assessed for system performance and stability. For each analyte, inter-day repeatability was obtained by calculating the RSD values for four calibration curve samples that were analyzed in different days, see SI Table S5. These four samples had identical concentration and came from the four sets of calibration curves that were evenly distributed throughout the run (see study design). For calculation of inter-day repeatability RSD values, two concentrations between LLOQ and ULOQ were included for each metabolite, as presented in Table S5. The inter-day repeatability of each metabolite is shown in Table S5 as the RSD values for analyte response. The RSD values for inter-day repeatability were found to be between 1.7% and 26.6% and generally lower than 20% for most part of the analytes, in agreement to previous observations ^16^. Gln and Glu shown interday repeatability higher than 20% due to a decreased peak intensity for their internal standard. In order to correct this and improve repeatability, we suggest a two-times increase in internal standard concentration for Gln and Glu in the final method, see supporting information Table S2.

Another strategy to evaluate the feasibility of an analytical method is to quantify the precision of the method through coefficients of variation (CV) or relative standard deviation (RSD) of quality control samples ^43,44^. We calculated RSD values for each metabolite quantified in the 36 QC pooled samples distributed throughout 600 injections and compare these values to the study variation (RSD in study samples), see Table S5. Among the 34 investigated metabolites, 28 presented RSD values below 30% which is the threshold for analytical reproducibility using intra-study QC samples ^3^. On the other hand, samples from the study-cohorts show high RSD values that were found to be from 2 to 158 times higher than the RSD values for pooled QC samples, thus demonstrating a considerably higher level of biological variation than the level of technical variation. The comparison between these variations is graphically captured by Figure 3, where violin plots representing the distribution of metabolite concentrations in pools and cohort samples are shown. As observed in Figure 3, the variation of each of the metabolites quantified in the QC pooled samples was lower in comparison to the variation in the samples of the study-cohort. It is important to highlight that the six analytes that presented RSD values higher than 30% in the pooled samples, IndS, AADA, N-MNA, Gly, ADMA, HCit, were found to have concentrations under the quantification limit. Moreover, some of these metabolites, for example N-MNA, were not detected in most of the pooled samples. These observations demonstrate that the method is sensitive and reproducible for the quantification of the 28 metabolites: GBB, GCDCA, GDCA, Trp, Ile, TUDCA, AzelA, CA, Leu, GUDCA, Ala, TDCA, TCDCA, Phe, Glu, GCA, Cit, TCA, Taurine, Tyr, LCA, CDCA, DCA, □-OHB, Gln, Kynu, UDCA, GLCA.

**Figure 3.**
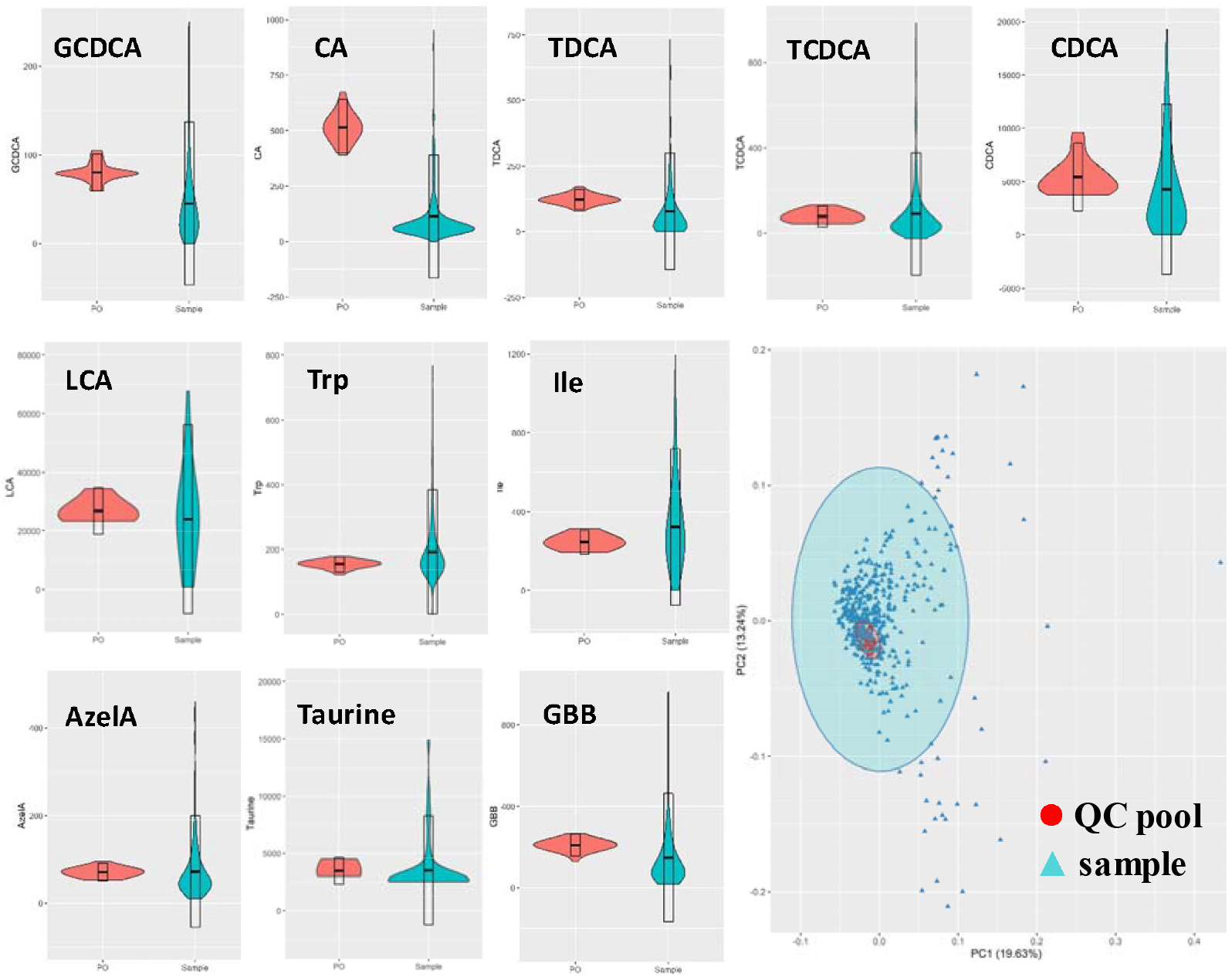
Violin plots for distribution of metabolite concentrations across human fecal samples representing a cohort with different degrees of liver disease (n=475) and fecal pooled samples (PO), used as replicates (n=36) for quality control. Bottom right: PCA scores plot for fecal metabolites in pooled samples and samples from the investigated cohort.

While RSD values are useful for evaluating each metabolite individually, another way to represent the total variation of the QC samples in relation to the samples of the cohort is to obtain a principal component analysis (PCA) plot, shown in Figure 3. As observed in Figure 3 and SI Figure S3, QC pooled samples were found to cluster closely to each other in the center of the PCA, whereas study samples were spread in the PCA plot. As the PCA plot of Figure 3 projected the contribution of all metabolites for the variations of the groups, the overall variation of the QC pooled samples was concluded to be low, indicating that is possible to obtain high quality data from the method described in the present study.

The data obtained for QC pooled samples can be further used for correcting systematic variations across analytical batches and to align and postprocess the full dataset before starting the statistical analysis that will be used for answering the clinical question ^43,44^. An important conclusion from the results, is that the high variation of the samples represents biological variation and not technical variation, providing evidence that our method can give insights into relevant biological processes.

Herein, we demonstrated the feasibility of our method for a large-scale study. Moreover, we report average concentrations (μmol g^-1^ of dried sample) of the metabolites quantified by our method in feces from healthy controls, see Table 1.

Our method allowed us to quantify 12 bile acids together with 10 amino acids and azelaic acid, gamma-butyrobetaine and beta-hydroxybutyrate in fecal samples from healthy controls. The metabolites UDCA, GUDCA, glutamine, glycine, homocitruline, aminoadipic acid, indoxyl sulphate and N-MNA presented concentrations below the quantification limit in fecal samples from healthy controls. However, these compounds were found in fecal samples from other cohort that represents patients with alcoholic liver disease. The average concentrations of bile acids obtained in fecal samples from healthy controls were comparable to the values reported by Shafaei et al., 2021, in a study using LC-MS quantitative method for determination of 12 bile acids ^7^. The concentrations of the secondary bile acids obtained in the present study were also found to be in the same range of values reported in other studies using HPLC-MS ^24^. These findings corroborate the reproducibility and feasibility of our method.

Finally, we calculated ratios between secondary to primary bile acids in feces as they represent a way of quantifying the conversion of primary bile acids into secondary bile acids by the intestinal microbial flora. Some studies have been using these ratios to investigate how changes in gut microbial composition impact bile acid metabolism and how they are linked to liver diseases such as cirrhosis ^24,42^. The ratios between secondary to primary bile acids are reported to be typically lower in cirrhotic patients in comparison to healthy controls. In Table 1, we report the ratios of secondary/primary bile acids in feces from healthy controls aiming for a future comparison with cohorts that represent patients with history of harmful drinking. Investigating the crosstalk between gut microbiota and liver through fecal metabolome provides a novel strategy for understanding the molecular mechanisms behind alcoholic liver disease ^45,46^.

## 4. CONCLUSION

We have developed a reproducible and robust method for preparing fecal samples for the quantification of 28 metabolites active on the gut-liver axis: GBB, GCDCA, GDCA, Trp, Ile, TUDCA, AzelA, CA, Leu, GUDCA, Ala, TDCA, TCDCA, Phe, Glu, GCA, Cit, TCA, Taurine, Tyr, LCA, CDCA, DCA, □-OHB, Gln, Kynu, UDCA, GLCA. An advantage of the proposed method is the quantification of analytes with different polarities in a single chromatographic run, in which will be practical in clinical and translational research. Moreover, our high-throughput method provides a tool for investigating gut-liver axis metabolism in liver-related diseases efficiently using a noninvasive collected sample. The investigation of bile acid metabolism using fecal metabolomics can be strategically used to uncover the underlying mechanisms behind gut-liver impairment.

## 5. ACKNOWLEDGMENT

This study was financed by the European Union’s Horizon 2020 research and innovation programme (grant number 668031) and it was supported by the Novo Nordisk Foundation through Challenge Grant “MicrobLiver” (grant number NNF15OC0016692). We thank the managers GALAXY and MicrobLiver consortia, respectively, Louise Skovborg Just and Lise Ryborg.

## ASSOCIATED CONTENT

### Supporting Information

The contents of supporting Information are described below.

MS parameters of the dynamic MRM method that was optimized from Ahonen et al., 2019 for detection of metabolites in fecal samples. Parameters include MRM transition, polarity, retention time, fragmentor voltage (V), collision energy (V) (Table S1)

MS parameters of the dynamic MRM method that was optimized from Ahonen et al., 2019 for detection of the internal standards in fecal samples. Parameters include MRM transition, polarity, retention time, fragmentor voltage (V), collision energy (V). Final concentration of internal standards (IS) in the ISTDmix in ng mL^-1^ and was determined for each compound based on their limits of quantification (Table S2)

Metabolites identified by GCGC-MS in fecal samples from healthy controls. Columns represent the classes of compounds that belong to fatty acid metabolism, amino acid metabolism, carbohydrate metabolites, TCA and others (Table S3)

Limit of detection (LOD); limit of quantification (LOQ); linearity (R2); linear regression parameters for calibration curves: (A) slope and (B) intercept with standard deviation (SD); range of linearity by UHPLC-MS analysis of bile acids and amino acids (Table S4)

Metabolite-wise relative standard deviation (RSD) in pooled samples (PO), study samples (“Sample”), estimated ratio between biological variation and technical variation (“Ratio”), inter-day repeatability for two different concentrations (conc. 1 and conc. 2) in the range of LLOQ and ULOQ (Table S5)

Study design for evaluation of method reproducibility in a UHPLC-MS targeted platform. Quality control (QC) samples including blanks, calibration solutions and pooled samples were distributed throughout 600 injections. Four calibration curves were placed at the beginning, middle and at the end of the analysis. Analysis started with 2 blanks, 3 pooled samples, a calibration curve with 11 levels of concentration and 1 blank. A unit composed by a blank, a pool and 16 clinical samples was repeated until the next calibration curve (Figure S1)

Analytical stability of a targeted UHPLC-MS platform for quantitative analysis of metabolites related to gut-liver axis. Time related variation of metabolite response measured in quality control pooled samples representing replicates and samples representing a cohort. Interval of variation in QC pooled samples is represented in grey (Figure S2).

Violin plots for distribution of metabolite concentrations across human fecal samples representing a cohort with different degrees of liver disease (n=475) and fecal pooled samples (PO), used as replicates (n=36) for quality control (Figure S3).

## AUTHOR INFORMATION

### Present Addresses

† Copenhagen Prospective Studies on Asthma in Childhood, Herlev and Gentofte Hospital, University of Copenhagen, 1165 Copenhagen, Denmark.

### Author Contributions

A.Z., M.T., T.H., A.K. and C.LQ. conceptualized the study. A.Z., M.K. and C.L.Q. designed the research. A.Z., A.W., M.K., A.F.B, K.Z., and I.M.M. performed the experiments. A.Z., T.S., A.W. and M.A. analyzed the data. T.S. and M.A. developed data analysis software and algorithms. M.T., T.H. and A.K. provided patient samples from their clinical study. A.Z. and C.L.Q. wrote the paper. A.Z., T.S., A.W., M.A. and C.L.Q. revised the paper.

### Notes

The authors declare no competing financial interest.

## REFERENCES

(1) Vuckovic, D. Current Trends and Challenges in Sample Preparation for Global Metabolomics Using Liquid Chromatography-Mass Spectrometry. Anal. Bioanal. Chem. 2012, 403 (6), 1523–1548.

(2) Karu, N.; Deng, L.; Slae, M.; Guo, A. C.; Sajed, T.; Huynh, H.; Wine, E.; Wishart, D. S. A Review on Human Fecal Metabolomics: Methods, Applications and the Human Fecal Metabolome Database. Anal. Chim. Acta 2018, 1030, 1–24.

(3) Broadhurst, D.; Goodacre, R.; Reinke, S. N.; Kuligowski, J.; Wilson, I. D.; Lewis, M. R.; Dunn, W. B. Guidelines and Considerations for the Use of System Suitability and Quality Control Samples in Mass Spectrometry Assays Applied in Untargeted Clinical Metabolomic Studies. Metabolomics 2018, 14 (6).

(4) Theodoridis, G. A.; Gika, H. G.; Want, E. J.; Wilson, I. D. Liquid Chromatography-Mass Spectrometry Based Global Metabolite Profiling: A Review. Anal. Chim. Acta 2012, 711 (January), 7–16.

(5) Trošt, K.; Ahonen, L.; Suvitaival, T.; Christiansen, N.; Nielsen, T.; Thiele, M.; Jacobsen, S.; Krag, A.; Rossing, P.; Hansen, T.; Dragsted, L. O.; Legido-Quigley, C. Describing the Fecal Metabolome in Cryogenically Collected Samples from Healthy Participants. Sci. Rep. 2020, 10 (1), 1–8.

(6) Hosseinkhani, F.; Dubbelman, A.-C.; Karu, N.; Harms, A. C.; Hankemeier, T. Towards Standards for Human Fecal Sample Preparation in Targeted and Untargeted LC-HRMS Studies. Metabolites 2021, 11 (6), 364.

(7) Shafaei, A.; Rees, J.; Christophersen, C. T.; Devine, A.; Broadhurst, D.; Boyce, M. C. Extraction and Quantitative Determination of Bile Acids in Feces. Anal. Chim. Acta 2021, 1150.

(8) Chen, M. X.; Wang, S. Y.; Kuo, C. H.; Tsai, I. L. Metabolome Analysis for Investigating Host-Gut Microbiota Interactions. Journal of the Formosan Medical Association. 2019, pp S10–S22.

(9) Krautkramer, K. A.; Fan, J. Gut Microbial Metabolites as Multi-Kingdom Intermediates. Nat. Rev. Microbiol. 2020, 19, 77–94.

(10) Deda, O.; Virgiliou, C.; Orfanidis, A.; Gika, H. G. Study of Fecal and Urinary Metabolite Perturbations Induced by Chronic Ethanol Treatment in Mice by UHPLC-MS/MS Targeted Profiling. Metabolites 2019, 9 (10).

(11) Ma, C.; Han, M.; Heinrich, B.; Fu, Q.; Zhang, Q.; Sandhu, M.; Agdashian, D.; Terabe, M.; Berzofsky, J. A.; Fako, V.; Ritz, T.; Longerich, T.; Theriot, C. M.; McCulloch, J. A.; Roy, S.; Yuan, W.; Thovarai, V.; Sen, S. K.; Ruchirawat, M.; Korangy, F.; Wang, X. W.; Trinchieri, G.; Greten, T. F. Gut Microbiome–Mediated Bile Acid Metabolism Regulates Liver Cancer via NKT Cells. Science. 2018.

(12) Caffaratti, C.; Plazy, C.; Mery, G.; Tidjani, A.-R.; Fiorini, F.; Thiroux, S.; Toussaint, B.; Hannani, D.; Le Gouellec, A. What We Know So Far about the Metabolite-Mediated Microbiota-Intestinal Immunity Dialogue and How to Hear the Sound of This Crosstalk. Metabolites 2021, 11 (6), 406.

(13) Arab, J. P.; Karpen, S. J.; Dawson, P. A.; Arrese, M.; Trauner, M. Bile Acids and Nonalcoholic Fatty Liver Disease: Molecular Insights and Therapeutic Perspectives. Hepatology 2017, 65 (1), 350–362.

(14) Lavelle, A.; Sokol, H. Gut Microbiota-Derived Metabolites as Key Actors in Inflammatory Bowel Disease. Nat. Rev. Gastroenterol. Hepatol. 2020, 17, 223–237.

(15) Salic, K.; Kleemann, R.; Wilkins-Port, C.; McNulty, J.; Verschuren, L.; Palmer, M. Apical Sodium-Dependent Bile Acid Transporter Inhibition with Volixibat Improves Metabolic Aspects and Components of Nonalcoholic Steatohepatitis in Ldlr-/-.Leiden Mice. PLoS One 2019, 14 (6), 1–22.

(16) Ahonen, L.; Jäntti, S.; Suvitaival, T.; Theilade, S.; Risz, C.; Kostiainen, R.; Rossing, P.; Orešič, M.; Hyötyläinen, T. Targeted Clinical Metabolite Profiling Platform for the Stratification of Diabetic Patients. Metabolites 2019, 9 (9), 1–19.

(17) Lake, A. D.; Novak, P.; Shipkova, P.; Aranibar, N.; Robertson, D. G.; Reily, M. D.; Lehman-Mckeeman, L. D.; Vaillancourt, R. R.; Cherrington, N. J. Branched Chain Amino Acid Metabolism Profiles in Progressive Human Nonalcoholic Fatty Liver Disease. Amino Acids 2015, 47 (3), 603–615.

(18) Cheng, S.; Wiklund, P.; Autio, R.; Borra, R.; Ojanen, X.; Xu, L.; Törmäkangas, T.; Alen, M. Adipose Tissue Dysfunction and Altered Systemic Amino Acid Metabolism Are Associated with Non-Alcoholic Fatty Liver Disease. PLoS One 2015, 10 (10), 1–17.

(19) Newgard, C. B.; An, J.; Bain, J. R.; Muehlbauer, M. J.; Stevens, R. D.; Lien, L. F.; Haqq, A. M.; Shah, S. H.; Arlotto, M.; Slentz, C. A.; Rochon, J.; Gallup, D.; Ilkayeva, O.; Wenner, B. R.; Yancy, W. S.; Eisenson, H.; Musante, G.; Surwit, R. S.; Millington, D. S.; Butler, M. D.; Svetkey, L. P. A Branched-Chain Amino Acid-Related Metabolic Signature That Differentiates Obese and Lean Humans and Contributes to Insulin Resistance. Cell Metab. 2009, 9 (4), 311–326.

(20) Anesi, A.; Rubert, J.; Oluwagbemigun, K.; Orozco-Ruiz, X.; Nöthlings, U.; Breteler, M. M. B.; Mattivi, F. Metabolic Profiling of Human Plasma and Urine, Targeting Tryptophan, Tyrosine and Branched Chain Amino Acid Pathways. Metabolites 2019, 9 (11).

(21) Clària, J.; Moreau, R.; Fenaille, F.; Amorós, A.; Junot, C.; Gronbaek, H.; Coenraad, M. J.; Pruvost, A.; Ghettas, A.; Chu-Van, E.; López-Vicario, C.; Oettl, K.; Caraceni, P.; Alessandria, C.; Trebicka, J.; Pavesi, M.; Deulofeu, C.; Albillos, A.; Gustot, T.; Welzel, T. M.; Fernández, J.; Stauber, R. E.; Saliba, F.; Butin, N.; Colsch, B.; Moreno, C.; Durand, F.; Nevens, F.; Bañares, R.; Benten, D.; Ginès, P.; Gerbes, A.; Jalan, R.; Angeli, P.; Bernardi, M.; Arroyo, V. Orchestration of Tryptophan-Kynurenine Pathway, Acute Decompensation, and Acute-on-Chronic Liver Failure in Cirrhosis. Hepatology 2019, 69 (4), 1686–1701.

(22) Muthulakshmi, S.; Saravanan, R. Efficacy of Azelaic Acid on Hepatic Key Enzymes of Carbohydrate Metabolism in High Fat Diet Induced Type 2 Diabetic Mice. Biochimie 2013, 95 (6), 1239–1244.

(23) Reiter, S.; Dunkel, A.; Metwaly, A.; Panes, J.; Haller, D.; Hofmann, T.; Berg, W. Development of a Highly-Sensitive UHPLC - MS / MS Quantitation Method for Fecal Bile Acids and Application on Crohn’ s Disease Studies. J. Agric. Food Chem. 2021, 69 (17), 5238–5251.

(24) Kakiyama, G.; Muto, A.; Takei, H.; Nittono, H.; Murai, T.; Kurosawa, T.; Hofmann, A. F.; Pandak, W. M.; Bajaj, J. S. A Simple and Accurate HPLC Method for Fecal Bile Acid Profile in Healthy and Cirrhotic Subjects: Validation by GC-MS and LC-MS. Journal of Lipid Research. 2014, pp 978–990.

(25) Northfield, T. C.; McColl, I. Postprandial Concentrations of Free and Conjugated Bile Acids down the Length of the Normal Human Small Intestine. Gut 1973, 14 (7), 513–518.

(26) Xie, G.; Wang, L.; Chen, T.; Zhou, K.; Zhang, Z.; Li, J.; Sun, B.; Guo, Y.; Wang, X.; Wang, Y.; Zhang, H.; Liu, P.; Nicholson, J. K.; Ge, W.; Jia, W. A Metabolite Array Technology for Precision Medicine. Anal. Chem. 2021, 93 (14), 5709–5717.

(27) Katsidzira, L.; Ocvirk, S.; Wilson, A.; Li, J.; Mahachi, C. B.; Soni, D.; DeLany, J.; Nicholson, J. K.; Zoetendal, E. G.; O’Keefe, S. J. D. Differences in Fecal Gut Microbiota, Short-Chain Fatty Acids and Bile Acids Link Colorectal Cancer Risk to Dietary Changes Associated with Urbanization Among Zimbabweans. Nutr. Cancer 2019, 71 (8), 1313–1324.

(28) Zheng, X.; Qiu, Y.; Zhong, W.; Baxter, S.; Su, M.; Li, Q.; Xie, G.; Ore, B. M.; Qiao, S.; Spencer, M. D.; Zeisel, S. H.; Zhou, Z.; Zhao, A.; Jia, W. A Targeted Metabolomic Protocol for Short-Chain Fatty Acids and Branched-Chain Amino Acids. Metabolomics 2013, 9 (4), 818–827.

(29) Røst, L. M.; Brekke Thorfinnsdottir, L.; Kumar, K.; Fuchino, K.; Eide Langørgen, I.; Bartosova, Z.; Kristiansen, K. A.; Bruheim, P. Absolute Quantification of the Central Carbon Metabolome in Eight Commonly Applied Prokaryotic and Eukaryotic Model Systems. Metabolites 2020, 10 (2), 74.

(30) Ismail, R.; Lee, H. Y.; Mahyudin, N. A.; Abu Bakar, F. Linearity Study on Detection and Quantification Limits for the Determination of Avermectins Using Linear Regression. Journal of Food and Drug Analysis. 2014, pp 407–412.

(31) Wickham, H. Ggplot 2: Elegant Graphics for Data Analysis; 2009; pp 10–1007.

(32) Yin, S.; Guo, P.; Hai, D.; Xu, L.; Shu, J.; Zhang, W.; Khan, M. I.; Kurland, I. J.; Qiu, Y.; Liu, Y. Optimization of GC/TOF MS Analysis Conditions for Assessing Host-Gut Microbiota Metabolic Interactions: Chinese Rhubarb Alters Fecal Aromatic Amino Acids and Phenol Metabolism. Anal. Chim. Acta 2017, 995, 21–33.

(33) Eijk, H. M. H. Van; Bloemen, J. G.; Dejong, C. H. C. Application of Liquid Chromatography – Mass Spectrometry to Measure Short Chain Fatty Acids in Blood. J. Chromatogr. B 2009, 877, 719–724.

(34) Zeng, M.; Cao, H. Fast Quantification of Short Chain Fatty Acids and Ketone Bodies by Liquid Chromatography-Tandem Mass Spectrometry after Facile Derivatization Coupled with Liquid-Liquid Extraction. J. Chromatogr. B Anal. Technol. Biomed. Life Sci. 2018, 1083 (January), 137–145.

(35) Orešic M, Anderson G, Mattila I, Manoucheri M, Soininen H, Hyötyläinen T, B. C. Targeted Serum Metabolite Profiling Identifies Metabolic Signatures in Patients with Alzheimer’ s Disease, Normal Pressure Hydrocephalus and Brain Tumor. Front. Neurosci. 2018, 11 (January), 1–7.

(36) Rodríguez-Coira, J.; Delgado-Dolset, M. I.; Obeso, D.; Dolores-Hernández, M.; Quintás, G.; Angulo, S.; Barber, D.; Carrillo, T.; Escribese, M. M.; Villaseñor, A. Troubleshooting in Large-Scale LC-ToF-MS Metabolomics Analysis: Solving Complex Issues in Big Cohorts. Metabolites 2019, 9 (11), 1–17.

(37) Krautbauer S, Bu□chler C, L. G. Relevance in the Use of Appropriate Internal Standards for Accurate Quantification Using LC–MS/MS: Tauro-Conjugated Bile Acids as an Example. Anal. Chem. 2016, 88 (22), 10957–10961.

(38) Xu, W.; Zhong, C.; Zou, C.; Wang, B.; Zhang, N. Analytical Methods for Amino Acid Determination in Organisms. 2020, 1071–1088.

(39) Cohen S.A.; D.P., M. Synthesis of a Fluorescent Derivatizing Reagent, 6-Aminoquinolyl-N-Hydroxysuccinimidyl Carbamate, and Its Application for the Analysis of Hydrolysate Amino Acids via High-Performance Liquid Chromato.Pdf. Anal. Biochem. 1993, 211 (2), 279–287.

(40) Song, Y.; Xu, C.; Kuroki, H.; Liao, Y.; Tsunoda, M. Recent Trends in Analytical Methods for the Determination of Amino Acids in Biological Samples. J. Pharm. Biomed. Anal. 2018, 147, 35–49.

(41) Lajczak-McGinley, N. K.; Porru, E.; Fallon, C. M.; Smyth, J.; Curley, C.; McCarron, P. A.; Tambuwala, M. M.; Roda, A.; Keely, S. J. The Secondary Bile Acids, Ursodeoxycholic Acid and Lithocholic Acid, Protect against Intestinal Inflammation by Inhibition of Epithelial Apoptosis. Physiol. Rep. 2020, 8 (12), 1–11.

(42) Kakiyama, G.; Pandak, W. M.; Gilleve, P. M.; Hylemon, P. B.; Heuman, D. M.; Daita, K.; Takei, H.; Muto, A.; Nittono, H.; Ridlon, J. M.; White, M. B.; Noble, N. A.; Monteith, P.; Fuchs, M.; Thacker, L. R.; Masoumeh; Bajaj, J. S. Modulation of the Fecal Bile Acid Profile by Gut Microbiota in Cirrhosis. J. Hepatol. 2013, 58 (5), 949–955.

(43) Godzien, J.; Alonso-Herranz, V.; Barbas, C.; Armitage, E. G. Controlling the Quality of Metabolomics Data: New Strategies to Get the Best out of the QC Sample. Metabolomics 2015, 11 (3), 518–528.

(44) Gika, H. G.; Theodoridis, G. A.; Earll, M.; Wilson, I. D. A QC Approach to the Determination of Day-to-Day Reproducibility and Robustness of LC-MS Methods for Global Metabolite Profiling in Metabonomics/Metabolomics. Bioanalysis 2012, 4 (18), 2239–2247.

(45) Szabo, G. Gut-Liver Axis in Alcoholic Liver Disease. Gastroenterology 2015, 148 (1), 10–36.

(46) Liang, Q.; Wang, C.; Li, B.; Zhang, A. Metabolomics of Alcoholic Liver Disease: A Clinical Discovery Study. RSC Adv. 2015, 5 (98), 80381–80387.

