## Supporting information for "High-throughput UHPLC-MS to screen metabolites in feces for gut metabolic health"

**Abstract:** (1) Background: Feces are the product of our diets and have been linked to diseases of the gut, including Chron’s disease and metabolic diseases such as diabetes. For screening metabolites in heterogeneous samples such as feces, it is necessary to use fast and reproducible analytical methods that maximize metabolite detection. (2) Methods: As sample preparation is crucial to obtain high quality data in MS-based clinical metabolomics, we developed a novel, efficient and robust method for preparing fecal samples for analysis with a focus in reducing aliquoting and detecting both polar and non-polar metabolites. Fecal samples (n= 475) from patients with alcohol-related liver disease and healthy controls were prepared according to the proposed method and analyzed in an UHPLC-QQQ targeted platform in order to obtain a quantitative profile of compounds that impact liver-gut axis metabolism. (3) Results: MS analyses of the prepared fecal samples have shown reproducibility and coverage of n=28 metabolites, mostly comprising bile acids and amino acids. We report metabolite-wise relative standard deviation (RSD) in quality control samples, inter-day repeatability, LOD, LOQ and range of linearity. The average concentrations for 135 healthy participants are reported here for clinical applications. (4) Conclusions: our high-throughput method provides an efficient tool for investigating gut-liver axis metabolism in liver-related diseases using a noninvasive collected sample.

**Keywords:** bile acids; fecal metabolomics; gut-liver axis; targeted metabolomics; sample preparation

Appendix A

EXPERIMENTAL SECTION

Sample collection

Stool samples were collected by the participants in their own home 24 hours prior to the scheduled visit to the research clinic. Participants were instructed to store the stool at −20 °C in their freezer immediately after sampling and, to transport the sample to the clinic in a cooling bag with ice. The stool samples were immediately stored in the research clinic at −80 °C upon arrival. Original fecal samples were sent to University of Copenhagen where they were cryogenically drilled to produce 200 mg aliquots that were handed over to Steno Diabetes Center Copenhagen. Fecal aliquots were then stored at −80 °C until further use.

Instrumentation

Initially, selected reaction monitoring (SRM) mode was used to determine the transitions between the precursor and fragment ions that produce the most intense fragments for each of the three additional analytes. SRM transitions of the other 31 analytes were defined as previously described by Ahonen et al., 2019 [16]. For each of the 34 determined SMR ion transitions, the following MS parameters were further optimized in order to ensure higher sensitive: fragmentor voltages, collision energies (CE), and cell accelerator voltages.

Preparation of pooled samples

After preparing the samples, the remaining supernatant was used for preparing pooled samples for quality control (QC) and the leftover was stored at -80 ˚C for further analyses. For this study, pooled samples were produced by mixing together all the 475 fecal supernatants produced in the step of derivatization. The resulting pooled supernatants were divided in aliquots of 20 μL which were prepared individually and identically to the original samples by adding 20 μL of ISTDmix following derivatization with AQC.

Study design for evaluation of analytical performance and suitability

Pooled samples representing replicates were included several times at the beginning of the analytical run and, also at intervals throughout the analysis to ensure system stability, see Figure S1. For quantification of metabolites, calibration curves were included at the beginning, at the middle and at the end of the analysis, see supporting information Figure S1, in order to evaluate and correct possible sensitivity loss following the progression of analysis. Each calibration level was spiked with ISTDmix and further derivatized with AQC, following dilution in methanol/water 1:1 (v/v) and preparation pattern identical to that of the original samples. The range of concentrations in the calibration curves was adapted from Ahonen et al., 2019 in order to fit the higher concentration of bile acids found in feces in relation to plasma [16].

**Table S 1.** MS parameters of the dynamic MRM method that was optimized from Ahonen et al., 2019 for detection of metabolites in fecal samples. Parameters include MRM transition, polarity, retention time, fragmentor voltage (V), collision energy (V).

| **Compound** | **Abbreviation** | **Molecular Weight (MW)** | **Retention time** | **MRM transition** | **Polarity** | **Fragmentor Voltage (V)** | **Collision Energy (V)** | **Cell Accelerator Voltage (V)** |
| --- | --- | --- | --- | --- | --- | --- | --- | --- |
| DL-2-Aminoadipic Acid | AADA | 161.2 | 2.6 | 330.2–160.1 | Negative | 150 | 10 | 1 |
| Asymmetric dimethylarginine | ADMA and SDMA | 202.3 | 2.4 | 371.2–201.2 * | Negative | 150 | 5 | 5 |
|  |  |  | 2.4 | 371.2–156.1 | Negative | 150 | 20 | 1 |
| L-Alanine | Ala | 89.1 | 2.6 | 258.1–88.1 | Negative | 100 | 15 | 3 |
| Azelaic acid | AzelA | 188.2 | 3.8 | 187.2–169 | Negative | 150 | 10 | 2 |
|  |  |  | 3.8 | 187.2–125.2 * | Negative | 150 | 15 | 2 |
| L-3-hydroxybutyric Acid | β-OHB | 104.1 | 0.7 | 103.2–59.2 | Negative | 100 | 5 | 1 |
| Cholic Acid | CA | 408.6 | 6.4 | 407.3–407.3 * | Negative | 250 | 10 | 3 |
|  |  |  | 6.4 | 407.3–343.3 | Negative | 250 | 35 | 1 |
| Deoxychenocholic Acid | CDCA | 392.6 | 7.0 | 391.3–391.3 | Negative | 200 | 10 | 4 |
| L-Citrulline | Cit | 175.2 | 2.5 | 344.4–174.2 | Negative | 150 | 4 | 7 |
| Deoxycholic Acid | DCA | 392.6 | 7.0 | 391.2–345.3 * | Negative | 200 | 35 | 4 |
|  |  |  | 7.0 | 391.2–327.2 | Negative | 200 | 40 | 4 |
| Gamma-butyrobetaine | GBB | 146.2 | 0.85 | 147.2–88.1 * | Positive | 100 | 16 | 1 |
|  |  |  | 0.85 | 147.2–60.2 | Positive | 100 | 13 | 1 |
| Glycocholic Acid | GCA | 465.6 | 5.95 | 464.3–402.1 | Negative | 250 | 40 | 4 |
|  |  |  | 5.95 | 464.3–74.1 * | Negative | 250 | 45 | 7 |
| Glycochenodeoxycholic Acid | GCDCA | 449.6 | 6.5 | 448.3–448.3 | Negative | 200 | 40 | 2 |
|  | GCDCA | 449.6 | 6.5 | 448.3–74.1 | Negative | 200 | 55 | 4 |
| Glycodeoxycholic Acid | GDCA | 449.6 | 6.6 | 448.3–74.2 | Negative | 200 | 55 | 4 |
| Glycolithocholic acid | GLCA | 433.3 | 7.1 | 432.3–388.1 | Negative | 200 | 25 | 4 |
|  |  |  |  | 432.3–74.1* | Negative | 200 | 45 | 4 |
| L-Glutamine | Gln | 146.1 | 2.4 | 315.3–145.1 | Negative | 100 | 9 | 6 |
| L-Glutamic Acid | Glu | 147.1 | 2.45 | 316.1–146.1 | Negative | 100 | 6 | 6 |
| Glycine | Gly | 75.1 | 2.4 | 244.1–74.1 | Negative | 200 | 7 | 4 |
| Glycoursodeoxycholic Acid | GUDCA | 449.6 | 6.0 | 448.3–386 | Negative | 250 | 40 | 2 |
|  |  |  | 6.0 | 448.3–74.1 * | Negative | 250 | 45 | 2 |
| L-Homocitrulline | HCit | 189.2 | 2.6 | 358.3–188.1 | Negative | 200 | 10 | 1 |
|  |  |  | 2.6 | 358.3–145 * | Negative | 150 | 25 | 2 |
| Indoxyl Sulfate | IndS | 213.2 | 2.55 | 212–132 * | Negative | 100 | 15 | 2 |
|  |  |  | 2.55 | 212–80 | Negative | 100 | 20 | 2 |
| L-Kynurenine | Kynu | 208.2 | 3.65 | 377–316.1 | Negative | 150 | 5 | 2 |
|  |  |  | 3.65 | 377–207 * | Negative | 150 | 5 | 5 |
| Lithocholic acid | LCA | 376.3 | 7.6 | 375.3–375.3 | Negative | 240 | 10 | 4 |
|  |  |  | 7.6 | 375.3–45 | Negative | 240 | 100 | 4 |
| L-Leucine | Leu and Ile | 131.2 | 3.72 | 300.2–130.2 | Negative | 100 | 10 | 1 |
| L-Isoleucine | Ile | 131.2 | 3.6 | 300.2–130.2 | Negative | 100 | 10 | 1 |
| N-methyl-nicotinamide | N-MNA | 136.2 | 1.05 | 137.1–108.1 | Positive | 100 | 15 | 2 |
|  |  |  |  | 137.1–80.2 * | Positive | 100 | 26 | 2 |
| L-Phenylalanine | Phe | 165.2 | 3.95 | 334.2–164 | Negative | 100 | 10 | 1 |
| Taurine | Taurine | 125.2 | 2.32 | 294.1–124.1 * | Negative | 100 | 10 | 2 |
|  |  |  |  | 294.1–80.1 | Negative | 100 | 55 | 2 |
| Taurocholic Acid | TCA | 515.7 | 5.7 | 514.3–123.8 | Negative | 300 | 65 | 5 |
|  |  |  | 5.7 | 514.3–80.2 * | Negative | 300 | 95 | 1 |
| Taurochenodeoxycholic Acid | TDCA and TCDCA | 499.3 | 6.05 | 498.3–107.1 | Negative | 250 | 80 | 4 |
|  |  |  | 6.15 | 498.3–80.1 * | Negative | 300 | 90 | 4 |
| Taurodeoxycholic Acid |  | 499.3 | 6.15 | 498.3–498.3 | Negative | 250 | 10 | 4 |
| L-Tryptophan | Trp | 204.2 | 4.27 | 373.2–203.1 | Negative | 150 | 7 | 2 |
| Tauroursodeoxycholic Acid | TUDCA | 499.7 | 5.7 | 498.3–107.1 | Negative | 250 | 65 | 5 |
|  |  |  |  | 498.3–80.1 * | Negative | 250 | 70 | 1 |
| L-Tyrosine | Tyr | 181.2 | 3.97 | 350.2–180.1 | Negative | 100 | 7 | 5 |
| Ursodeoxycholic acid | UDCA | 392.3 |  | 391.3–391.3 | Negative | 250 | 10 | 4 |
|  |  |  | 7.0 | 391.3–289.3 | Negative | 250 | 45 | 4 |

*ion transition used for quantification;

MRM transitions of amino acids represent the ion transition for the adduct with AQC reagent.

**Table S 2.** MS parameters of the dynamic MRM method that was optimized from Ahonen et al., 2019 for detection of the internal standards in fecal samples. Parameters include MRM transition, polarity, retention time, fragmentor voltage (V), collision energy (V). Final concentration of internal standards (IS) in the ISTDmix in ng mL^-1^ and was determined for each compound based on their limits of quantification.

| **Compound** | **IS concentration**  **(ng mL^-1^)** | **Molecular Weight (MW)** | **Retention time**  **(min)** | **MRM transition** | **Polarity** | **Fragmentor Voltage (V)** | **Collision Energy (V)** | **Cell Accelerator Voltage (V)** |
| --- | --- | --- | --- | --- | --- | --- | --- | --- |
| AADA-d3 | 30000 | 164.2 | 2.6 | 333.2–145.2 | Negative | 100 | 20 | 2 |
| ADMA-d7 | 10000 | 209.8 | 2.45 | 378–208.3 | Negative | 100 | 10 | 5 |
| Ala-d4 | 5000 | 93.1 | 2.6 | 262.1–92.1 | Negative | 100 | 5 | 6 |
| AzelA-d14 | 5000 | 202.3 | 3.7 | 201.2–137.2 | Negative | 150 | 10 | 2 |
| β-OHB-d4 | 100000 | 108.1 | 0.75 | 107.1–59.1 | Negative | 100 | 5 | 1 |
| CA-d4 | 500 | 412.3 | 6.37 | 411.3–411.3 | Negative | 250 | 10 | 3 |
| CDCA-d4 DCA-d4 | 500 | 396.6 | 7.0 | 395.2–395.2 | Negative | 300 | 10 | 4 |
| Cit-d4 | 1000 | 179.2 | 2.47 | 348.1–135.1 | Negative | 100 | 25 | 2 |
| GBB-d9 | 5000 | 154.7 | 0.87 | 155.2–87.3 | Positive | 100 | 13 | 1 |
| GCA-d4 | 250 | 469.6 | 5.97 | 468.3–74.1 | Negative | 250 | 45 | 1 |
| GCDCA-d4 GUDCA-d4 | 30000 | 453.6 | 6.55 | 452.3–74.1 | Negative | 250 | 40 | 1 |
| GUDCA-d4 | 10000 | 453.6 | 6.55 | 452.3–74.1 | Negative | 250 | 40 | 1 |
| GDCA-d6 | 30000 | 455.7 | 6.6 | 454.3–408.2 | Negative | 250 | 55 | 4 |
| GLCA-d4 | 1000 | 437.3 | 7.1 | 436.3–74.1 | Negative | 200 | 50 | 4 |
| Gln-d5 | 60000 | 151.2 | 2.46 | 320.1–150.1 | Negative | 100 | 5 | 1 |
| Glu-d5 | 10000 | 152.1 | 2.46 | 321.1–151.1 | Negative | 100 | 5 | 1 |
| Gly-13C,d2 | 10000 | 78.1 | 2.4 | 247–77.1 | Negative | 100 | 5 | 7 |
| HCit-2H4 | 10000 | 193.2 | 2.6 | 362.2–192.2 | Negative | 100 | 5 | 6 |
| IndS-d4 | 10000 | 217.3 | 2.55 | 216–136.1 | Negative | 100 | 15 | 2 |
| Kynu-13C6 | 30000 | 214.2 | 3.6 | 383.1–195.8 | Negative | 100 | 10 | 6 |
| LCA-d4 | 1000 | 380.3 | 7.55 | 379.3–379.3 | Negative | 240 | 100 | 4 |
| Leu-d10 Ile-d10 | 5000 | 141.2 | 3.6  3.55 | 310.1–140 | Negative | 125 | 10 | 2 |
| N-MNA-d4 | 750 | 140.2 | 1.3 | 141.2–84.2 | Positive | 100 | 20 | 7 |
| Phe-d5 | 1000 | 170.2 | 3.9 | 339.1–169.1 | Negative | 150 | 5 | 1 |
| Taurine-d4 | 1000 | 129.2 | 2.32 | 298.3–128.2 | Negative | 100 | 10 | 3 |
| TCA-d4 | 500 | 519.7 | 5.7 | 518.3–80 | Negative | 340 | 100 | 7 |
| TCDCA-d9 | 500 | 508.3 | 6.1 | 507.4–80.1 | Negative | 300 | 95 | 4 |
| Trp-d8 | 5000 | 212.3 | 4.25 | 381.2–211.2 | Negative | 100 | 10 | 5 |
| TUDCA-d4 | 250 | 503.7 | 5.7 | 502.3–80.1 | Negative | 250 | 70 | 1 |
| Tyr-d7 | 5000 | 188.2 | 2.97 | 357.1–187.2 | Negative | 100 | 10 | 1 |
| UDCA-d4 | 250 | 396.6 | 7.0 | 395.3–395.3 | Negative | 250 | 0 | 4 |

MRM transitions of amino acids represent the ion transition for the adduct with AQC reagent.

**Table S 3.** Metabolites identified by GCGC-MS in fecal samples from healthy controls. Columns represent the classes of compounds that belong to fatty acid metabolism, amino acid metabolism, carbohydrate metabolites, TCA and others.

| **Free fatty acids** | |  | **Amino acids** |  | **Carbohydrates** |  | **TCA metabolites /others** |
| --- | --- | --- | --- | --- | --- | --- | --- |
| Stearic acid | Nonanedioic acid |  | Methionine |  | Myo inositol |  | Citric acid |
| Oleic acid | Tetradecanoic acid |  | Serine |  | d-Ribose |  | Fumaric acid |
| Palmitic acid | 1,4-Butanediol |  | Glycine |  | L-(-)-Fucose |  | Succinic acid |
| Palmitoleic acid | 1-Octadecanol |  | Leucine |  | Sorbitol |  | Malic acid |
| 1-Monopalmitin | 1-Dodecanol |  | Phenylalanine |  | Maltose |  | Lactic acid |
| Linoleic acid | 3,4-Dihydroxybutanoic acid |  | Alanine |  | d-Galactose |  | 3-Indoleacetic acid |
| Tetradecanoic acid | 2,4-Dihydroxybutanoic acid |  | Proline |  | D-Arabinose |  | 3-Hydroxyphenylacetic acid |
| Butanoic acid | 4-Hydroxybutanoic acid |  | Valine |  | Lactic acid |  | Deoxycholic acid (bile acid) |
| Hexanoic acid | Benzoic acid |  | a-alanine |  | D-(-)-Fructofuranose |  |  |
| Pentanoic acid | Triethylene glycol |  | cadaverine |  | a -D-Glucopyranose |  |  |
| Heptanoic acid | Diethylene glycol |  |  |  | D-(+)-Galactopyranose |  |  |
| Octanoic acid | Glyceric acid |  |  |  | alpha-ketoglutaric acid |  |  |
| Nonanoic acid | Glycerol |  |  |  |  |  |  |
| Decanoic acid | Glycerol monostearate |  |  |  |  |  |  |
| Heptadecanoic acid | 5-Aminovaleric acid |  |  |  |  |  |  |
| Pentadecanoic acid | 3-Methyladipic acid |  |  |  |  |  |  |

**Table S 4.** Limit of detection (LOD); limit of quantification (LOQ); linearity (R2); linear regression parameters for calibration curves: (A) slope and (B) intercept with standard deviation (SD); range of linearity by UHPLC-MS analysis of bile acids and amino acids.

| **Compound** | **LOD**  **(ng mL^-1^)** | **LOQ**  **(ng mL^-1^)** | **R^2^** | **slope** | **SD of slope** | **intercept** | **SD of intercept** | **LLOQ** | | **ULOQ** |
| --- | --- | --- | --- | --- | --- | --- | --- | --- | --- | --- |
| AADA | 253.87 | 769.30 | 0.951 | 0.00221 | 3.93 10^-6^ | 15.26658 | 0.170031 | 1000 | 50000 | |
| ADMA | 311.35 | 943.49 | 0.983 | 0.000184 | 5.11 10^-7^ | -0.11094 | 0.017319 | 1000 | 50000 | |
| Ala | 0.03 | 0.084 | 0.978 | 26.20151 | 0.082371 | -3.63255 | 0.221374 | 2 | 50000 | |
| AZelA | 139.47 | 422.64 | 0.91 | 0.000597 | 2.12 10^-6^ | -0.50651 | 0.025227 | 500 | 25000 | |
| β-OHB | 177.23 | 537.06 | 0.993 | 8.04 10^-6^ | 1.34 10^-8^ | 6.2210^-5^ | 0.000432 | 750 | 10000 | |
| CA | 70.78 | 214.49 | 0.999 | 0.00257 | 2.05 10^-6^ | -1.00791 | 0.055114 | 10 | 100000 | |
| CDCA | 0.69 | 2.09 | 0.998 | 0.004932 | 2.12 10^-6^ | 0.022947 | 0.001031 | 10 | 50000 | |
| Cit | 217.05 | 657.71 | 0.992 | 3.66 10^-5^ | 7.11 10^-8^ | -0.05226 | 0.002409 | 750 | 50000 | |
| DCA | 7.915 | 23.98 | 0.995 | 0.000124 | 7.69 10^-7^ | 0.000553 | 0.000296 | 25 | 100000 | |
| GBB | 153.43 | 464.92 | 0.992 | 0.000186 | 3.22 10^-7^ | 0.024853 | 0.008633 | 500 | 50000 | |
| GCA | 12.36 | 37.44 | 0.999 | 0.001841 | 2.20 10^-7^ | -0.19432 | 0.006893 | 50 | 100000 | |
| GCDCA | 3.81 | 11.54 | 0.969 | 0.015198 | 3.29 10^-5^ | 0.056013 | 0.017533 | 50 | 50000 | |
| GDCA | 12.71 | 38.51 | 0.989 | 0.009236 | 8.67 10^-5^ | -0.03423 | 0.035568 | 50 | 50000 | |
| GLCA | 6.36 | 19.29 | 0.997 | 0.005836 | 2.74 10^-5^ | -0.00354 | 0.011255 | 50 | 100000 | |
| Gln | 230.60 | 698.78 | 0.995 | 0.001883 | 8.47 10^-6^ | 1.57648 | 0.131603 | 750 | 25000 | |
| Glu | 174.43 | 528.58 | 0.969 | 0.000264 | 4.18 10^-7^ | 0.162696 | 0.01398 | 500 | 50000 | |
| Gly | 265.13 | 803.42 | 0.979 | 0.001627 | 4.96 10^-6^ | -2.15256 | 0.130738 | 1000 | 100000 | |
| GUDCA | 5.90 | 17.88 | 0.999 | 0.000256 | 1.28 10^-8^ | -0.05641 | 0.000458 | 50 | 100000 | |
| HCit | 295.98 | 896.90 | 0.985 | 0.000598 | 1.58 10^-6^ | -0.88806 | 0.053648 | 1000 | 50000 | |
| Ile | 32.20 | 97.57 | 0.944 | 0.0016 | 3.27 10^-5^ | -0.01189 | 0.015609 | 100 | 50000 | |
| IndS | 145.05 | 439.54 | 0.996 | 3.09 10^-5^ | 4.30 10^-8^ | -0.02029 | 0.001358 | 500 | 100000 | |
| Kynu | 171.43 | 519.47 | 0.972 | 0.481562 | 7.42 10^-4^ | -1045.78 | 25.0159 | 500 | 10000 | |
| LCA | 7.09 | 21.49 | 0.998 | 0.00247 | 1.11 10^-5^ | 0.016653 | 0.005307 | 10 | 100000 | |
| Leu | 0.02 | 0.05 | 0.972 | 45.95384 | 0.16 | -3.75385 | 0.250482 | 2 | 100000 | |
| N-MNA | 11.29 | 34.23 | 0.994 | 0.000535 | 4.23 10^-6^ | -0.00196 | 0.001832 | 100 | 50000 | |
| Phe | 34.75 | 105.31 | 0.941 | 0.00034 | 7.50 10^-6^ | -0.0028 | 0.003578 | 250 | 50000 | |
| Taurine | 81.68 | 247.51 | 0.995 | 0.000618 | 3.44 10^-7^ | -1.45427 | 0.015284 | 250 | 50000 | |
| TCA | 37.39 | 113.32 | 0.999 | 0.004373 | 1.16 10^-6^ | 0.615186 | 0.049558 | 100 | 100000 | |
| TCDCA | 0.07 | 0.21 | 0.998 | 0.033012 | 4.10 10^-6^ | -0.02961 | 0.000682 | 50 | 100000 | |
| TDCA | 23.52 | 71.27 | 0.966 | 0.026576 | 0.000437 | -0.12875 | 0.189402 | 100 | 100000 | |
| Trp | 225.02 | 681.88 | 0.932 | 0.00255 | 9.95 10^-6^ | -6.27553 | 0.173872 | 750 | 100000 | |
| Tyr | 165.60 | 501.82 | 0.992 | 0.000319 | 4.92 10^-7^ | -0.34754 | 0.016009 | 500 | 100000 | |
| UDCA | 5.65 | 17.12 | 0.997 | 0.004278 | 1.90 10^-5^ | 0.008358 | 0.007324 | 50 | 50000 | |
| TUDCA | 60.80 | 184.23 | 0.997 | 0.003023 | 1.30 10^-6^ | -0.31566 | 0.05569 | 500 | 50000 | |

**Table S 5.** Metabolite-wise relative standard deviation (RSD) in pooled samples (PO), study samples (“sample”), estimated ratio between biological variation and technical variation (“ratio”), inter-day repeatability for two different concentrations (conc. 1 and conc. 2) in the range of LLOQ and ULOQ.

| **Compound** | **PO** | **Sample** | **Ratio** | **Inter-day repeatability (n=4)** | | |
| --- | --- | --- | --- | --- | --- | --- |
|  |  |  |  | **conc. 1** | **conc. 2** | |
| GBB | 12.9 | 2.03 10^3^ | 158 | 4.2 | | 2.2 |
| GCDCA | 13 | 1.73 10^3^ | 133 | 9.9 | | 15.3 |
| GDCA | 29 | 2.05 10^3^ | 70.8 | 15.2 | | 7.7 |
| Trp | 7.68 | 491 | 64 | 5.9 | | 3.8 |
| Ile | 12.2 | 411 | 33.5 | 12.9 | | 8.5 |
| TUDCA | 21.3 | 708 | 33.2 | 16.8 | | 3.55 |
| AzelA | 14.4 | 477 | 33.2 | 7.1 | | 19.8 |
| CA | 12.2 | 362 | 29.7 | 3.8 | | 2.1 |
| Leu | 16.9 | 481 | 28.4 | 2.8 | | 11.5 |
| GUDCA | 13.4 | 314 | 23.5 | 3.7 | | 1.6 |
| Ala | 16.1 | 230 | 14.2 | 17.6 | | 6.5 |
| TDCA | 15.7 | 247 | 15.7 | 9.7 | | 21.4 |
| Phe | 11 | 170 | 15.6 | 3.6 | | 1.9 |
| TCDCA | 20.5 | 304 | 14.8 | 4.2 | | 12.5 |
| Glu | 16.2 | 219 | 13.5 | 26.6 | | 10.6 |
| GCA | 22.9 | 300 | 13.1 | 3.2 | | 1.7 |
| Cit | 14.9 | 99.6 | 6.68 | 2.3 | | 8.5 |
| TCA | 13.2 | 65.8 | 5 | 4.1 | | 8.2 |
| Taurine | 12.2 | 322 | 26. 3 | 2.5 | | 5.6 |
| Tyr | 6.24 | 25.5 | 4.09 | 4.4 | | 5.1 |
| LCA | 15.9 | 105 | 6.65 | 3.2 | | 5.6 |
| CDCA | 29.1 | 100 | 3.44 | 3.4 | | 2.6 |
| DCA | 9.79 | 88 | 8.89 | 6.4 | | 1.1 |
| b-OHB | 21.5 | 273 | 11.8 | 5.0 | | 13.8 |
| Gln | 28.9 | 108 | 3.0 | 11.5 | | 26.0 |
| IndS | 128 | 182 | 1.42 | 6.5 | | 20.0 |
| Kynu | 23.4 | 31 | 1.33 | 6.2 | | 3.7 |
| AADA | 80.8 | 410 | 5.08 | 8.8 | | 18.3 |
| UDCA | 24.0 | 116 | 4.84 | 12.8 | | 4.3 |
| GLCA | 18.5 | 23.9 | 1.29 | 2.1 | | 10.5 |
| Gly^a^ | - | - | - | 16.7 | | 16.5 |
| N-MNA | 41.7 | 99.2 | 2.38 | 16.9 | | 4.7 |
| ADMA | 34.6 | 43.1 | 1.24 | 2.5 | | 5.2 |
| HCit | 86.4 | 104 | 1.21 | 6.4 | | 4.9 |

^a^ non-detected in 50% of samples

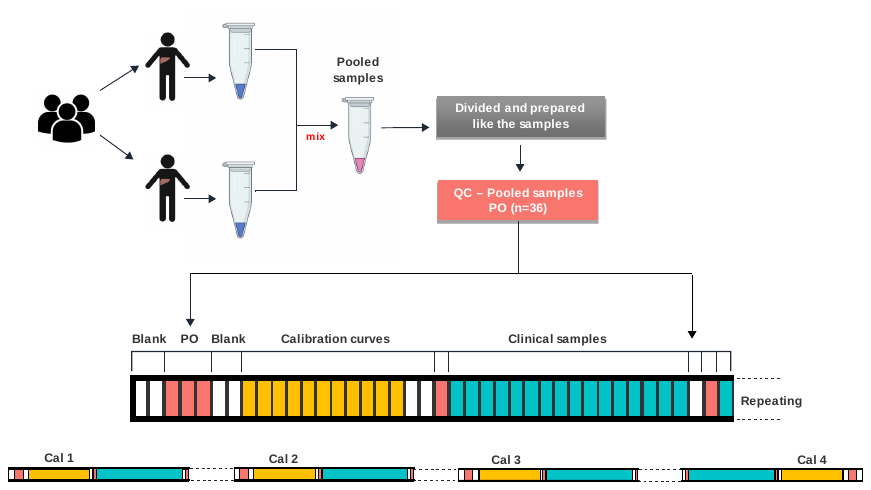

**Figure S 1.** Study design for evaluation of method reproducibility in a UHPLC-MS targeted platform. Quality control (QC) samples including blanks, calibration solutions and pooled samples were distributed throughout 600 injections. Four calibration curves were placed at the beginning, middle and at the end of the analysis. Analysis started with 2 blanks, 3 pooled samples, a calibration curve with 11 levels of concentration and 1 blank. A unit composed by a blank, a pool and 16 clinical samples was repeated until the next calibration curve.

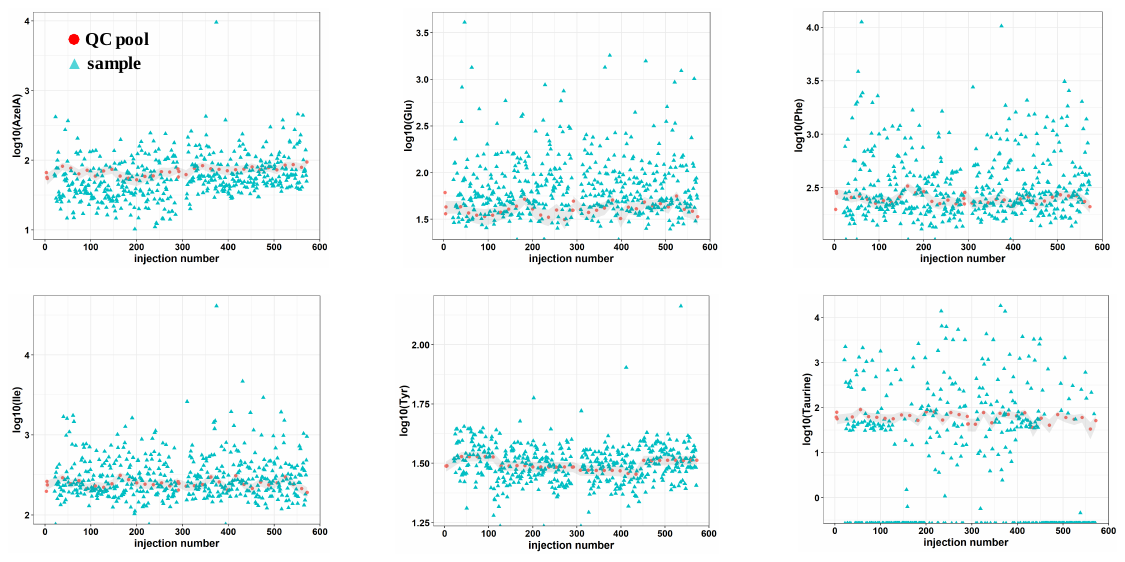

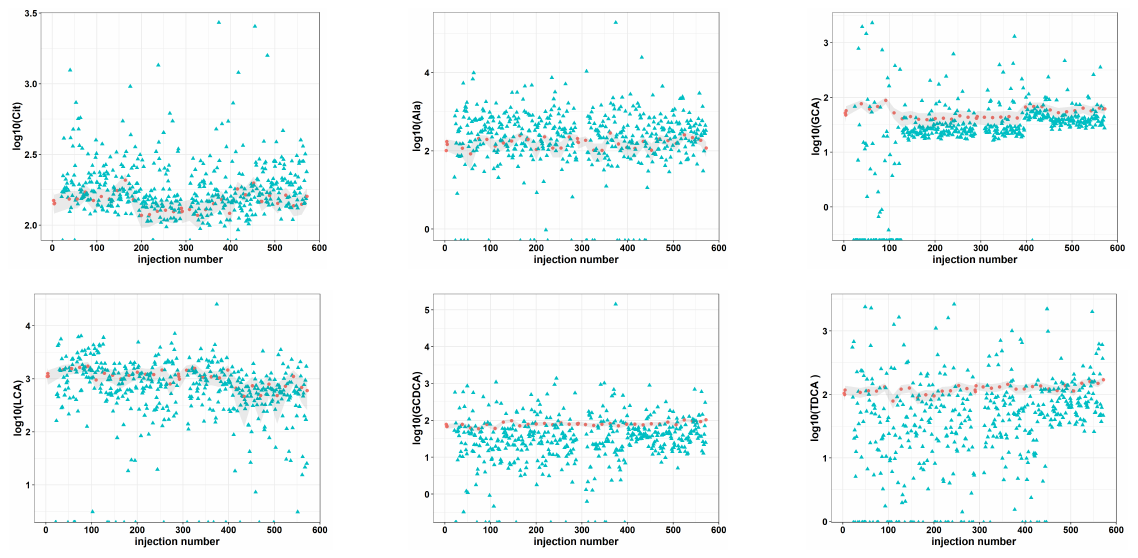

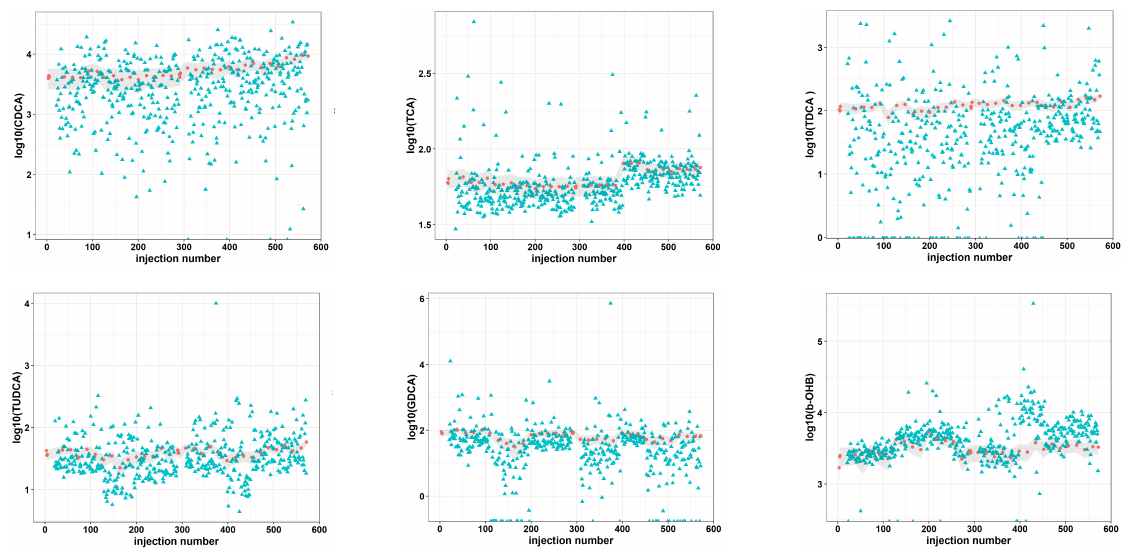

**Figure S 2.** Analytical stability of a targeted UHPLC-MS platform for quantitative analysis of metabolites related to gut-liver axis. Time related variation of metabolite response measured in quality control pooled samples representing replicates and samples representing a cohort. Interval of variation in QC pooled samples is represented in grey.

**Figure S 3.** Violin plots for distribution of metabolite concentrations across human fecal samples representing a cohort with different degrees of liver disease (n=475) and fecal pooled samples (PO), used as replicates (n=36) for quality control.
